# Intestinal commensal bacteria promote *Bactrocera dorsalis* larval development through vitamin B6 synthesis pathway

**DOI:** 10.1101/2024.04.15.589666

**Authors:** Jian Gu, Zhichao Yao, Bruno Lemaitre, Zhaohui Cai, Hongyu Zhang, Xiaoxue Li

## Abstract

**Background:** The gut microbiota can facilitate host growth under nutrient-constrained conditions. However, whether this effect is limited to certain bacterial species remains largely unclear, and the relevant mechanisms remain to be thoroughly investigated.

**Results:** Here, we found that the microbiota was required for *Bactrocera dorsalis* larval growth under poor diet conditions. Mono-association experiments revealed that *Enterobacteriaceae* and some *Lactobacilli* promoted larval growth. Of 27 tested bacterial strains, 15 strains significantly promoted larval development, and the *Enterobacteriaceae cloacae* N29 isolate exhibited the most obvious promoting effect. Bacterial genome-wide association study (GWAS) revealed that the vitamin B6 synthesis pathway was critical for *E. cloacae* growth promotion. The deletion of *pdxA* responsible for the vitamin B6 biosynthesis deprived the mutant strains of larval growth promotion function, indicating that *pdxA* gene was crucial for promoting larval growth in the N29 strain. Importantly, supplementation of vitamin B6 to poor diet successfully rescued the axenic larval growth phenotype of *B. dorsalis*.

**Conclusion:** Our results suggest that gut microbes promote insect larval growth by providing vitamin B6 under nutrient scarcity in *B. dorsalis*.

## 1. Introduction

Insects, like other animals, are closely associated with microorganisms. The microorganisms have great effect on insects’ physiological, ecological, and evolutionary functions [1,2]. Microorganisms, including bacteria, archaea, fungi, viruses, and others, may have enduring or transient interrelationships with their insect hosts, which may be beneficial or detrimental to insect health [1]. Although microorganisms may be pathogenic to insects, thus reducing host viability and increasing morbidity [3], commensal microorganisms within insects could be helpful or even essential for the host’s survival.

Commensal microorganisms provide protection to the host against various stresses, such as temperature, drought, and heavy metal stresses, as well as biological stresses from nematodes, fungi, and parasites [4]. The greater wax moth, *Galleria mellonella,* could digest polyethylene with the aid of intestinal microbiome [5]. The endosymbiont *Buchnera* increases the thermotolerance of its host aphid, and this effect is associated with the *ibpA* gene in aphid [6]. The bacterium *Hamiltonella defensa* can provide protection against parasitoids in pea aphids [7–11]. Another symbiont *Regiella insecticola* can offer protection against fungal infection and enhance resistance to parasitoids, thus promoting pea aphid survival [12,13]. This protection is also common in other insects. Gut symbiotic bacterium *Burkholderia* increases the resistance of its host bean bug *Riptortus pedestris* against pesticide fenitrothion [14]. In *Oryzaephilus surinamensis* insect, symbionts affect cuticle thickness, thus promoting desiccation resistance under dry conditions [15]. Two core microbes *Snodgrassella alvi* and *Lactobacillus bombicola* in bumble bee increase host’s survival, when host is exposed to selenate [16]. The bacterial communities in eggs and larvae of chironomids play a role in protecting hosts from toxicants [17]. These findings illustrate the contribution of microbiota to host fitness by supplementing host metabolic pathways or increasing resistance of hosts to stresses [1].

Many insects live in nutrient-limited environments, which significantly impacts insect physiological activity. Under these nutrient-constrained conditions, insect microbes can provide nutrients to their hosts. A case in point is that aphids live by sucking plant phloem sap, but the essential amino acid tryptophan is not present in phloem sap [18,19], and this amino acid absent in phloem sap is supplied by the aphid’s primary symbiotic bacterium, *Buchnera aphidicola* [20–23]. Another example is that the nutrient supply of glassy-winged leafhopper *Homalodisca coagulate* is dependent on two symbiotic bacteria *Baumannia cicadellinicola* and *Sulcia muelleri*, *of which B.cicadellinicola* provides the host with vitamins and coenzymes, while *S. muelleri* provides some essential amino acids, thus ensuring the metabolic functions of host [24]. In Hemiptera insect feeding on bast sap, symbiotic bacterium *Sulcia sp.* has evolved a dual symbiotic relationship with host and a variety of endosymbiotic bacteria, and the reduced genome of *Sulcia sp.* makes it dependent on other symbiotic bacteria to synthesize all essential amino acids [25,26]. Symbiotic bacterium

*Candidatus Erwinia dacicola* in olive fruit fly *Bactrocera oleae* can extract nitrogen from environmental wastes such as bird droppings, thus providing the adult fly with essential amino acids and increasing their reproductive capacity [27]. Many studies have highlighted the role of the microbiota in providing the host with choline, thiamine, and amino acids in model organism *Drosophila melanogaster* [28–30]. Several studies have indicated that the presence of symbiotic bacteria promotes the growth of *Drosophila* larvae [31–33]. An integrated nutritional network established based on a systematic nutritional screening reveals the complex interactions between the host *Drosophila* and its microbiota species [34].

Previous research focuses on the identification of a single microbiota species altering host traits. However, insect microbiomes are often composed of various bacteria from different orders and families, working jointly to affect the host. Here we analyzed how the gut microbiomes of the invasive horticultural pest *Bactrocera dorsalis* collectively function in altering or promoting host adaptability. *B. dorsalis* is a fruit fly damaging over 350 different fruits and vegetables, and microbiota in *B. dorsalis* plays multiple roles in the host physiological processes including cold adaptation, survival after irradiation, reproductive behavior, and invasion [35–38].

This study mainly investigated the microbiota’s role in larval development of *B. dorsalis*. 16s rRNA sequencing revealed that as larvae grew and developed, the richness and diversity of the gut microbial communities of *B. dorsalis* gradually decreased. Microbiota was required for fly growth under a poor-nutrition diet conditions. Mono-association experiment indicated that of 27 tested bacterial species, at least over 15 different bacterial species promoted larval growth on the poor diet. The bacterial genome-wide association study (GWAS) demonstrated that 23 pathways in bacteria were positively associated with host larval growth (indicated by body length). Further analysis demonstrated that vitamin B6 synthesis pathway in the symbiotic bacteria was the key pathway regulating larval growth.

## Results

### Microbial diversity decreases during *B. dorsalis* larval development

We first investigated the gut microbiota diversity of 3-instar larvae of *B. dorsalis* including the 1^st^-3^rd^ instars (Figure S1)[39]. We performed 16S rRNA sequencing of the samples of these 3 different instars collected from the lab. The results showed that the number of OTUs shared by all three instars was 700 out of the total 2002 OTUs, indicating that about 35% of the OTUs were relatively stable in the gut of larvae of 3 different instars, and the gut of the first instar had the larger number of OTUs than the other 2 instars (Figure 1A). The results of UniFrac non-metric multidimensional scaling (NMDS) analysis also showed that there were differences in intestinal microbial communities among 3 different instars. The unweighted UniFrac NMDS analysis revealed that the gut microbiota of the first instar larvae and the third instar larvae belonged to two different clusters (Figure 1B). The weighted UniFrac NMDS analysis suggested that the first and second instar samples were separated, but the third instar samples were not clearly distinguished from the other two (Figure 1C). The above results jointly indicated that low-abundance microbial communities were instar-specific. However, the high-abundance species tended to be found in all three instars, suggesting a relatively high stability.

**Figure 1.**
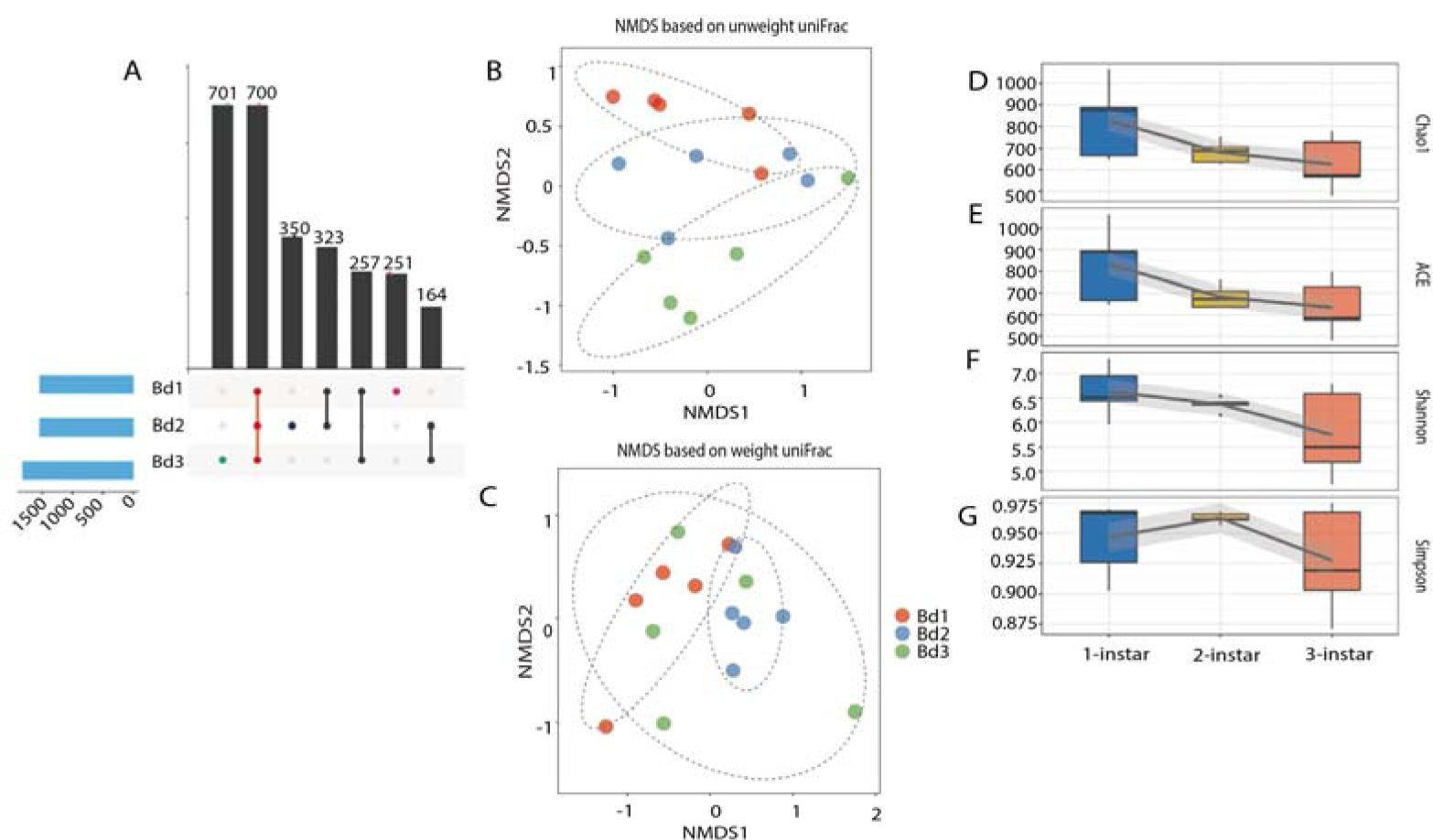
Diversity analysis of intestinal microbial community in larvae of 3 different instars. (A) Distribution of the common OTUs shared by 3 different instars and the unique OTUs specific to certain instar. (B) NMDS analysis based on unweighted UniFrac distance. (C) NMDS analysis based on weighted UniFrac distance. (D) Chao1 diversity index. (E) ACE diversity index. (F) Shannon diversity index. (G) Simpson diversity index. Bd1, Instar 1 larva; Bd 2, Instar 2 larva; Bd 3, Instar 3 larva.

Next, we analyzed the alpha diversity of the gut microbial communities of larvae in three different instars. The results showed that there was no significant difference in the Chao1 index, ACE index, Shannon index, and Simpson index of intestinal microbial communities among three different instars. However, the average value of all four indexes showed a clear decreasing trend with the larval development (Figure 1D-G). These results indicated that the richness and diversity of the gut microbiota were gradually decreased during larval development.

### Composition of gut microbial community in the larvae of *B. dorsalis*

Further, we examined the composition of gut microbial community in the larvae of *B. dorsalis*, and found that the larval gut microbiota was mainly composed of Proteobacteria, Firmicutes, Bacteroidetes, and Actinobacteria at the phylum level (Figure 2A). We also analyzed the relative abundance of bacteria at the family level. The results showed that Enterobacteriaceae and Streptococcaceae exhibited a high relative abundance in the larval gut at all three instars (Instar 1, I1; Instar 2, I2; and Instar 3, I3) indicating their stability (Figure 2B). Further analysis showed that the relative abundance of Enterobacteriaceae was significantly higher in the second instar larvae (52.70%± 2.541) than in the first instar larvae (28.55% ±3.716) (Figure 2C).

**Figure 2.**
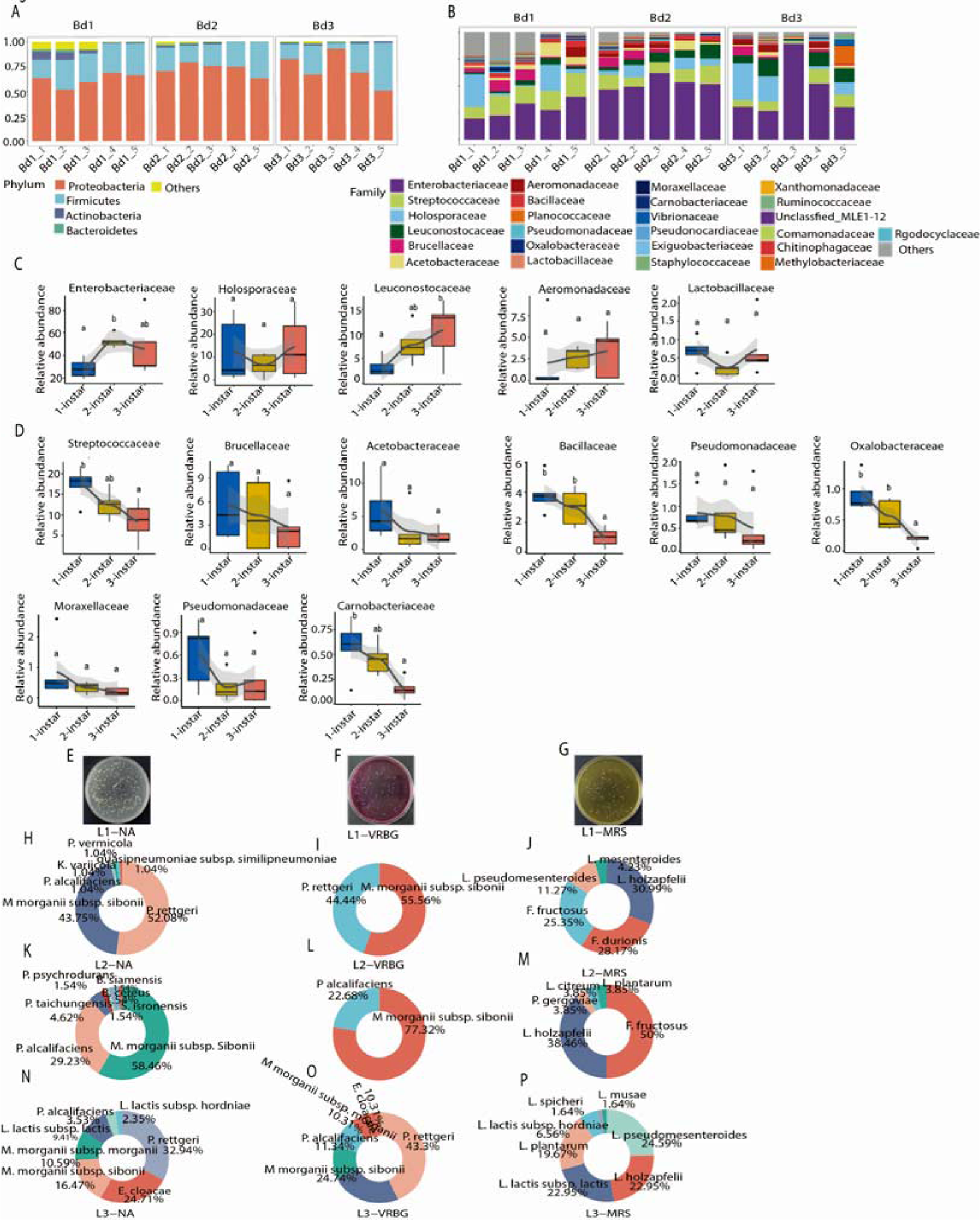
Composition of gut culturable microbiota in different instar larvae. (A) Gut microbiota composition at phylum level in different instar larvae. (B) Gut microbiota composition at family level in different instar larvae. Bd1_1, Bd1_2, Bd1_3, Bd1_4, and Bd1_5 represent the 5 biological replicates of instar-1 larvea, respectively; Bd2_1, Bd2_2, Bd2_3, Bd2_4, and Bd2_5 indicate the 5 biological replicates of instar-2 larvea, respectively; Bd3_1, Bd3_2, Bd3_3, Bd3_4, and Bd3_5 denote the 5 biological replicates of instar-3 larvea. (C) Relative abundance increase with larval instars. (D) Relative abundance decrease with larval instars. (E) Colony morphology on NA medium. (F) Colony morphology on VRBG medium. (G) Colony morphology on MRS medium. (H) Gut culturable microbiota composition of instar-1 larvae on NA medium. (I) Gut culturable microbiota composition of instar-1 larvae on VRBG medium. (J) Gut culturable microbiota composition of instar-1 larvae on MRS medium. (K) Gut culturable microbiota composition of instar-2 larvae on NA medium. (L) Gut culturable microbiota composition of instar-2 larvae on VRBG medium. (M) Gut culturable microbiota composition of instar-2 larvae on MRS medium. (N) Gut culturable microbiota composition of instar-3 larvae on NA medium. (O) Gut culturable microbiota composition of instar-3 larvae on VRBG medium. (P) Gut culturable microbiota composition of instar-3 larvae on MRS medium.

The relative abundance of Leuconostocaceae in the gut also increased significantly during development, with its relative abundance in the third instar larvae significantly higher than that in the first instar larvae (Figure 2C).

On the contrary, some microbiota families exhibited a clear decreasing trend with larval development. The relative abundances of Streptococcaceae, Bacillaceae, Oxalobacteraceae, and Carnobacteriaceae families were significantly lower in the I3 larvae than in the I1 larvae (Figure 2D). The relative abundances of Brucellaceae, Acetobacteracea, Pseudomonadaceae, Moraxellaceae and Pseudonocardiaceae were not statistically different in the gut during larval development, but still showed a clear decreasing trend (Figure 2D). In general, our results indicated that Enterobacteriaceae and Leuconostocaceae were stable in *B. dorsalis* with high relative abundance during larval development, suggesting their important role in larval growth and development of *B. dorsalis*.

### Composition of cultivable bacterial communities in the larvae of *B. dorsalis*

To further verify the above results, we cultured and isolated cultivable bacterial species using NA (a non-selective medium), VRBG (a selective medium for *Enterobacteriaceae*), and MRS (a selective medium for *Lactobacilli*) (Figure 2E-G). We obtained 246 isolated strains from NA medium (I1: 96, I2:65, and I3:85), 266 strains from VRBG medium (I1:72, I2:97, and I3:97), and 158 strains were identified on MRS Medium (I1:71, I2:26, and I3:61) (Table S1). In order to monitor changes in intestinal bacterial composition at different instars, we monitored bacterial counts on a specific bacteria medium plate. In the I1-NA, we identified *Providencia rettgeri* and *Morganella morganii subsp. sibonii* accounting for 52.08% and 43.75% (Figure 2H), respectively. In the I2-NA, *M. morganii subsp. sibonii* was a relatively high-abundance species (58.46%) (Figure 2K). Notably, the abundance of *Providencia alcalifaciens* increased from 1.04% (at I1) to 29.23% (at I2). In the I3-NA, *P. rettgeri* accounted for 32.94%; *E. cloacae* accounted for 24.71%; and *M. morganii subsp. sibonii* accounted for 24.71% (Figure 2N). These findings were in accordance with alpha diversity analysis results that high-abundance species were relatively stable, while low-abundance species were less stable.

We observed similar results in the VRBG medium. In the I1-VRBG, we identified *M. morganii subsp. sibonii* (55.56%) and *P. rettgeri* (44.44%) (Figure 2I). In the I2-VRBG, we observed *M. morganii subsp. sibonii* (77.73%) and *P. alcalifaciens* (22.68%) (Figure 2L). In the L3-VRBG, we found *P. rettgeri* (43.30%), *M. morganii subsp. Sibonii* (24.74%), and *P. alcalifaciens* (11.34%) (Figure 2O). Collectively, these data indicated that Enterobacteriaceae was relatively stable during *B. dorsalis* larval development, suggesting their important role in mediating larval growth.

Since *Lactobacilli* plays a crucial role in *Drosophila* and other animals’ growth [31–34], we cultured and isolated Lactobacillales using MRS selective medium, a selective medium for *Lactobacilli and* other species in Lactobacillales. In the I1-MRS, *Leuconostoc holzapfelii* accounted for 30.99%; *Fructobacillus durionis* accounted for 28.17%; *Leuconostoc pseudomesenteroides* accounted for 11.27%; and *Fructobacillus fructosus* and *L. pseudomesenteroides* accounted for 25.35% and 11.27% (Figure 2J). We also observed the change in *Lactobacillus* bacterial composition in I2 larvae (Figure 2M), and found that *F. fructosus* and *L. holzapfelii* were two dominant species, accounting for more than 88% of the total culturable *Lactobacillius* bacteria. In I3-MRS medium (Figure 2P), *L. pseudomesenteroides* accounted for 24.59%, and *L. holzapfelii* accounted for 22.95%. In general, *Providencia* and *Morganella* belonging to the *Enterobacteriaceaer* and *Leuconostoc* belonging to the *Leuconostocaceae* were the dominant cultivable bacteria, stably present in the gut of *B. dorsalis* larvae. In summary, the investigation of intestinal bacterial abundance of *B. dorsalis* larvae revealed that *Enterobacteriaceae* and *Leuconostoc* might play an important role in the growth and development of the *B. dorsalis* larvae.

### Single bacterium replenishment promotes larval growth

We raised axenic larvae (AX group) and conventional larvae (non-axenic larvae, CK group) on the artificial diet with 0.4% yeast concentrations. The results showed that larvae of the AX group exhibited a significantly shorter body length than the CK group feeding on the diet without yeast. Replenishment with culturable microbiota (AXA) partially rescued axenic larvae body length (Figure 3A).

**Figure 3.**
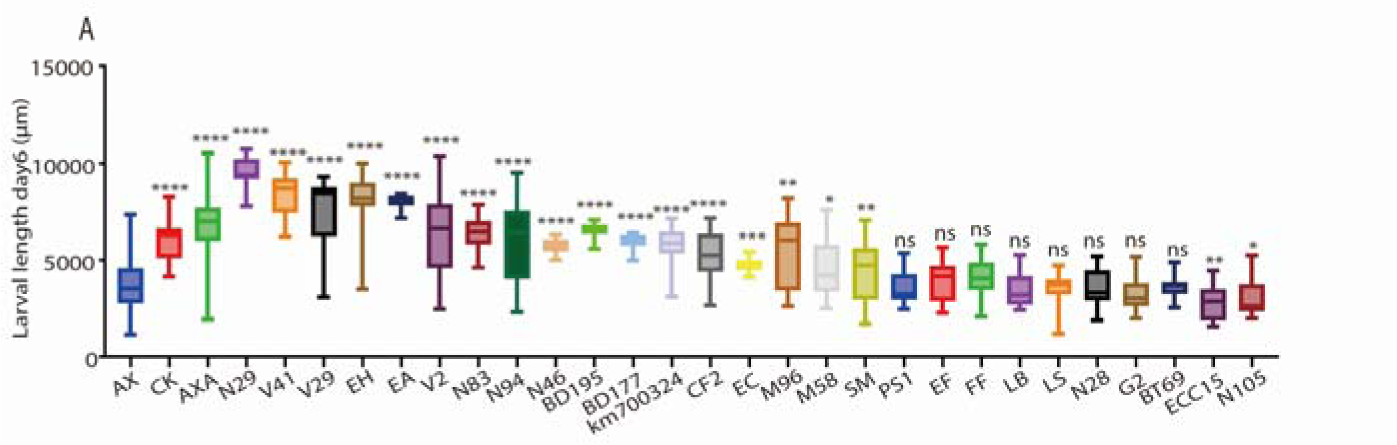
Effects of 27 bacterial strains mono-associated with larvae on larval growth. (A) The 27 strains which were mono-associated with larvae are BD177, BD195, BT69, CF2, EA, EC, ECC15, EF, EH, FF, G2, km700324, LB, LS, M58, M96, N105, N28, N29, N46, N83, N94, PS1, SM, V2, V29, and V41, respectively. AXA, Axenic larvae on a diet added with the total gut microbiota; AX, Axenic larvae; CK, control group (untreated wild larvae). The difference in mean body length of larvae were compared between treatment groups and AX group using Student’s t test. *, P < 0.05; **, P < 0.01; ****, P < 0.0001; ns, not statistically significant.

Next, we performed mono-association experiments to decipher the mechanism by which the microbiota promoted larval growth. We selected a total 27 bacterial strains, including 21 *B. dorsalis* gut bacterial strains (isolated in this study and those from lab stock), and 6 strains from *B. dorsalis* habitat (previous lab stock) (Table 1). The larvae were raised on 0.4% yeast diet, and their length were measured at day 6. The results showed that the average larval length of AX group was 3.79±0.10 mm, and that of AXA group was 6.71±0.16 mm, which was significantly higher than that of AX group. Mono-association experiment revealed that 15 strains out of 27 tested strains significantly promoted AX larval body length (Figure 3A). Among these 15 strains, AX larvae mono-associated with *E. cloacae* (N29), *M. morganii* (V41), *P. alcalifaciens* (V29), and *E. hormaechei* (EH) (larvae feeding on diets treated with one single strain) were significantly longer than AXA larvae replenished with total gut bacteria, indicating that these strains had a great impact on larval growth (Figure 3E). *E. cloacae* (N29) had the most prominent effect, the AX larvae mono-associated with this bacterium displayed more than twice the length of AX group without strain replenishment. Moreover, the symbiotic bacteria *Lactobacillis* including *L. plantarum* (M96) and *L. citreum* (M58) significantly promoted the growth of larvae, and the non-symbiotic *Escherichia coli* (EC) also significantly promoted the growth of larvae. Pathogenic bacteria *Serratia marcescens* (PS1) and *Providencia stuartii* (BT69) had no significant effect on larval growth, whereas *Psychrobacillus soli* (N105) and pathogenic *Erwinia carotovora Ecc15* significantly inhibited the growth of larvae. In general, all the *Enterobacteriaceae* strains and most *Lactobacillus* strains isolated from the intestine of *B. dorsalis* promoted larval growth, while other symbiotic bacteria have no effect on larval growth, and pathogenic bacteria had no or negative impact on larval growth.

**Table 1.** Candidate bacterium strains of mono-associated axenic larvae.

| Strain | Resource | Classification level |
| --- | --- | --- |
| BD177 | *Bactrocera dorsalis* | *Klebsiella michiganensis* |
| BD195 | *Bactrocera dorsalis* | *Citrobacter koseri* |
| BT69 | *Zelugodacus tau* | *Providencia stuartii* |
| CF2 | *Zelugodacus tau* | *Citrobacter portucalensis* |
| EA | *Bactrocera dorsalis* | *Enterobacter soli* |
| EC | DH5α strain | *Escherichia coli* |
| *Ecc15* | *Drosophila melanogaster* | *Erwinia carotovora* |
| EF | *Bactrocera dorsalis* | *Enterococcus faecalis* |
| EH | *Bactrocera dorsalis* | *Enterobacter hormaechei* |
| FF | *Bactrocera dorsalis* | *Fructobacillus fructosus* |
| G2 | *Bactrocera dorsalis* | *Agrobacterium fabrum* |
| km700324 | American Type Culture Collection | *Klebsiella michiganensis* |
| LB | *Bactrocera dorsalis* | *Lactobacillus brevis* |
| LS | *Bactrocera dorsalis* | *Lactobacillus spicheri* |
| M58 | *Bactrocera dorsalis* | *Leuconostoc citreum* |
| M96 | *Bactrocera dorsalis* | *Lactobacillus plantarum* |
| N105 | *Bactrocera dorsalis* | *Psychrobacillus soli* |
| N28 | *Bactrocera dorsalis* | *Paenibacillus pabuli* |
| N29 | *Bactrocera dorsalis* | *Enterobacter cloacae* |
| N46 | *Bactrocera dorsalis* | *Klebsiella quasipneumoniae* |
| N83 | *Bactrocera dorsalis* | *Providencia stuartii* |
| N94 | *Bactrocera dorsalis* | *Morganella morganii* |
| PS1 | *Phyllotreta striolata* | *Serratia marcescens* |
| SM | *Bactrocera dorsalis* | *Serratia nematodiphila* |
| V2 | *Bactrocera dorsalis* | *Providencia rettgeri* |
| V29 | *Bactrocera dorsalis* | *Providencia alcalifaciens* |
| V41 | *Bactrocera dorsalis* | *Morganella morganii* |

### Genome-wide association study of intestinal bacterial strains of *B. dorsalis*

To investigate the effects of the bacteria on host larval growth, we performed genome sequencing of these 27 strains. The phylogenetic tree constructed by the maximum likelihood method showed that *Klebsiella*, *Providencia*, *Morganella*, *Citrobacter*, *Serratia*, *Erwinia,* and *Escherichia* were close to each other, and they belonged to *Enterobacteriaceae* (Figure 4A). *Psychrobacillus*, *Paenibacillus,* and *Agrobacterium* were clustered together, and they belonged to the *lactobacilli*, lactic acid-producing bacteria, but they were far from the *Enterobacteriaceae*. The above evolutionary relationships might explain the reasons for differences in their ability to promote larval growth of *B. dorsalis*.

**Figure 4.**
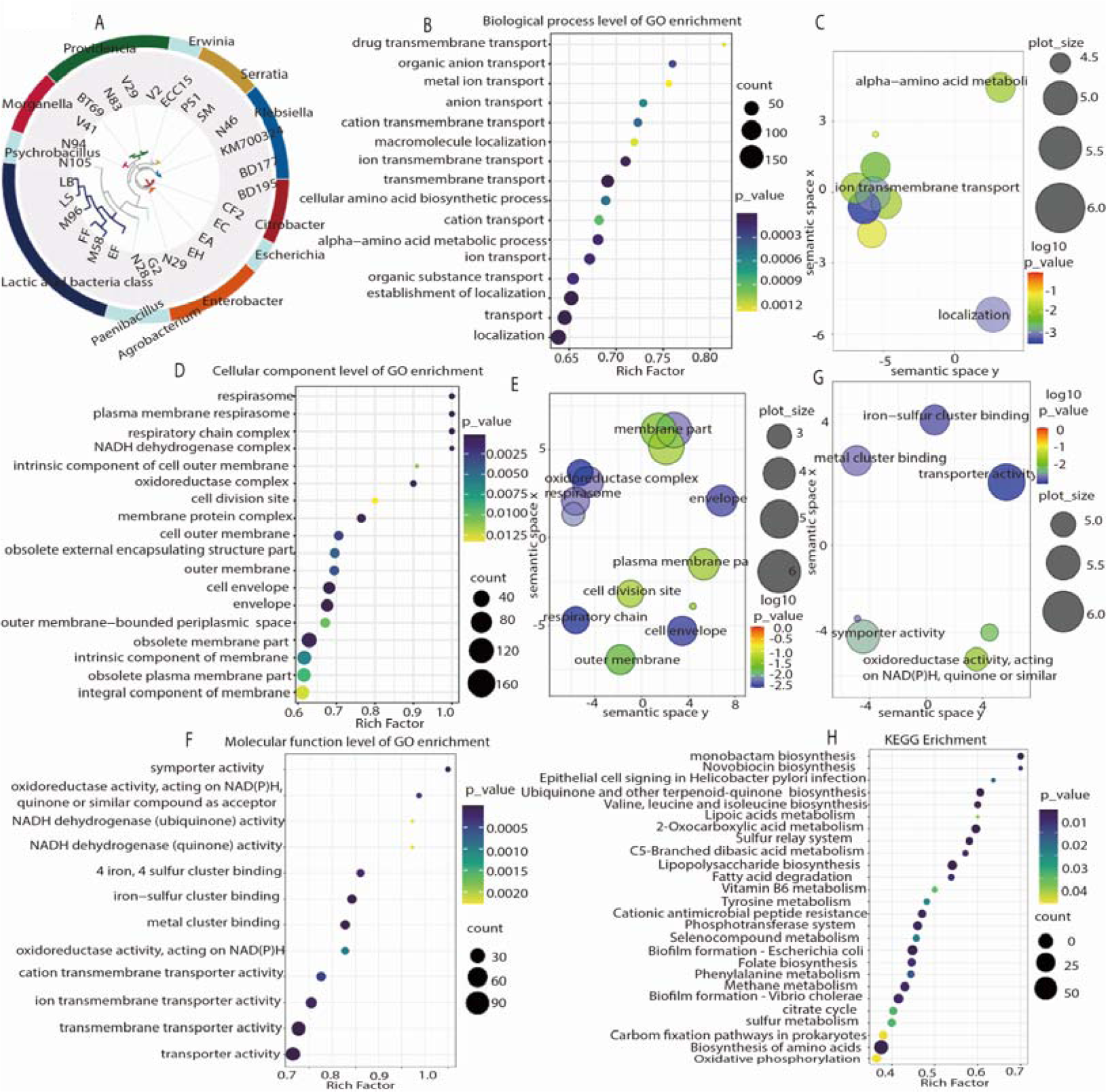
Genome-wide association study (GWAS) of intestinal bacteria in *B. dorsalis* larvae. (A) Phylogenetic tree of 27 strains. A total of 1,904 single copy genes in the 27 strain genomes were concatenated, and the total length of amino acid sequence was 15,742. (B-C) GO enrichment analysis of biological processes in which 3,417 larval growth-associated orthologous genes were enriched. Bubble map of 16 significantly enriched biological processes (B) and bubble map of 3 simplified biological process categories(C). (D-E) GO enrichment analysis of cellular components in which 3,417 larval growth-associated orthologous genes were enriched. Bubble map of 18 significantly enriched cellular components (D) and bubble map of 8 simplified cellular component categories. (F-G) GO enrichment analysis of molecular function in which 3,417 larval growth-associated orthologous genes were enriched. Bubble map of 12 significantly enriched molecular functions (F) and bubble map of 5 simplified molecular function categories.

Next, we analyzed the association between bacterial genome sequences and their larval growth promotion ability under a poor diet condition. Our bacterial genome-wide association study (GWAS) results showed that 5512 orthologous genes were significantly associated with body length. Among these genes, 3417 genes were positively associated with larval body length (Table 2). The GO enrichment analysis showed that these 3417 orthologous genes were mainly assigned to GO terms such as ion transport and binding, amino acid metabolism process, membrane structure composition, redox reaction, and respiratory chain, and these GO terms were mainly related to amino acids, micronutrition, and energy metabolism required for larval growth and development (Figure 4B-G).

**Table 2.** Information of Genome-wide association study of 27 strain genomes.

| Gene number | Orthologous gene (OG) | significantly associated OG | significantly positively associated OG |
| --- | --- | --- | --- |
| 110,940 | 9,567 | 5,512 | 3,417 |

KEGG pathway enrichment analysis results showed that all the 3417 genes were enriched in 23 pathways, such as tyrosine metabolism, Vitamin B6 synthesis, Valine, leucine, isoleucine biosynthesis, and folate biosynthesis (Figure 4H). The top five KEGG pathways with the largest gene numbers were amino acid biosynthesis pathway, biofilm formation-*Vibrio cholerae*, biofilm formation-*Escherichia coli*, lipopolysaccharide biosynthesis pathway, and oxidative phosphorylation pathway (Figure 4H).

### *E. cloacae* N29 promotes larvae growth by producing vitamin B6

Our data showed that *E. cloacae* N29 had the most significant promoting effect on larval growth under limited yeast diet conditions. Since yeast powders are rich in amino acids, vitamin B, and trace elements, we investigated 4 biosynthesis pathways including valine biosynthesis, leucine and isoleucine biosynthesis, vitamin B6 biosynthesis, and folate biosynthesis pathways in the *E. cloacae* N29. We selected 8 genes in *E. cloacae* N29 genome responsible for the above-mentioned 4 biosynthesis pathways, and we generated 8 *E. cloacae* N29 single mutants by deleting each of these genes (Table 3). To further reveal the roles of each of these four pathways in larval growth and development, we performed mono-association experiments of these 8 mutant strains. The results showed that axenic larvae on the diets supplemented with ΔYcgM, ΔtrpE, ΔpdxA2, ΔIlvA, ΔTdcB, ΔpabB, and ΔhpcE mutant strains exhibited successfully recovered body length (Figure 5A). However, supplementation with ΔpdxA mutant strains into axenic larval diet resulted in no significant difference in larval body length, indicating that *E. cloacae* N29 *pdxA* knockout strain lost the ability to promote larval growth (Figure 5A). Since the *pdxA* gene belonged to the vitamin B6 biosynthesis pathway, vitamin B6 biosynthesis played a important role in the promoting *B. dorsalis* larval growth.

**Figure 5.**
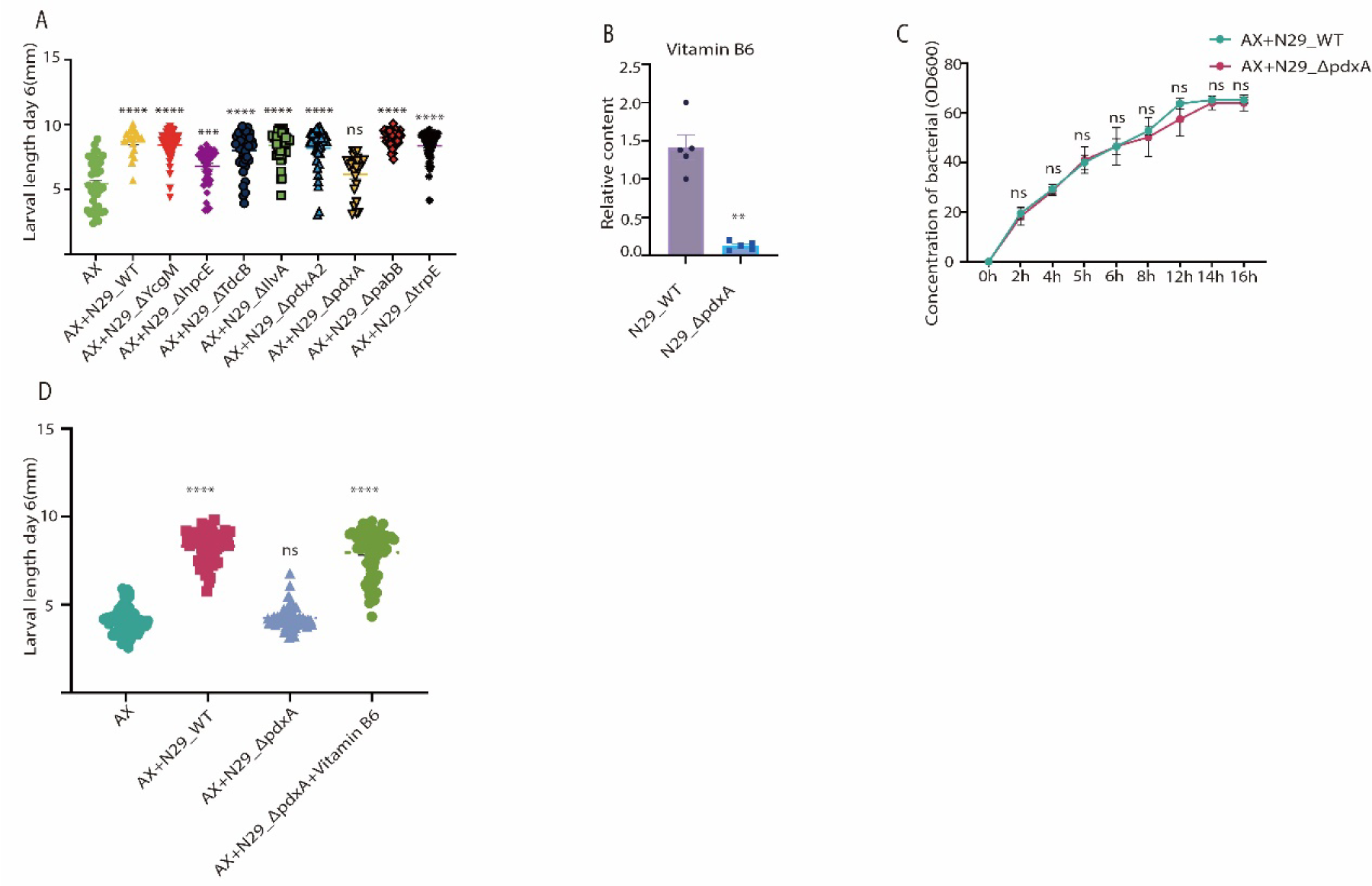
Effects of deletion of 8 genes in mutant strain of N29 on larvae growth. (A) Body length of 6-day-old larvae mono-associated with different N29 mutant strains. ΔTdcB, *TdcB*-deleted N29 mutant strain; Δ IlvA, *IlvA*-deleted N29mutant strain; Δ YcgM, *YcgM*-deleted N29 mutant strain; Δ hpcE, *hpcE*-deleted N29 mutant strain; Δ pdxA2, pdxA2-deleted N29 mutant strain; Δ pdxA,*pdxA*-deleted N29 mutant strain; Δ pabB, *pabB*-deleted N29 mutant; Δ trpE, *trpE*-deleted N29 mutant strain. (B) Relative vitamin B6 content of N29_WT and N29_Δ pdxA(5 biological replicates).(C) Growth curve of N29-WT and N29-Δ pdxA strain. (D) Body length of 6-day-old larvae in 4 groups of AX larvae, AX+N29_WT, AX+N29_Δ pdxA and AX+N29_Δ pdxA+ Vitamin B6. AX, axenic larvae. AX+N29_WT, axenic larvae on a diet added with wildtype N29; AX+N29_Δ pdxA, AX larvae on a diet added with *pdxA*-deleted N29 mutant strain; AX+N29_Δ pdxA+ Vitamin B6, AX larvae co-fed with *pdxA*-deleted N29 mutant and vitamin B6. The difference in mean body length of larvae were compared between treatment groups and AX group using Student’s t test. *, P < 0.05; **, P < 0.01; ****, P < 0.0001; ns, not statistically significant.

**Table 3.**
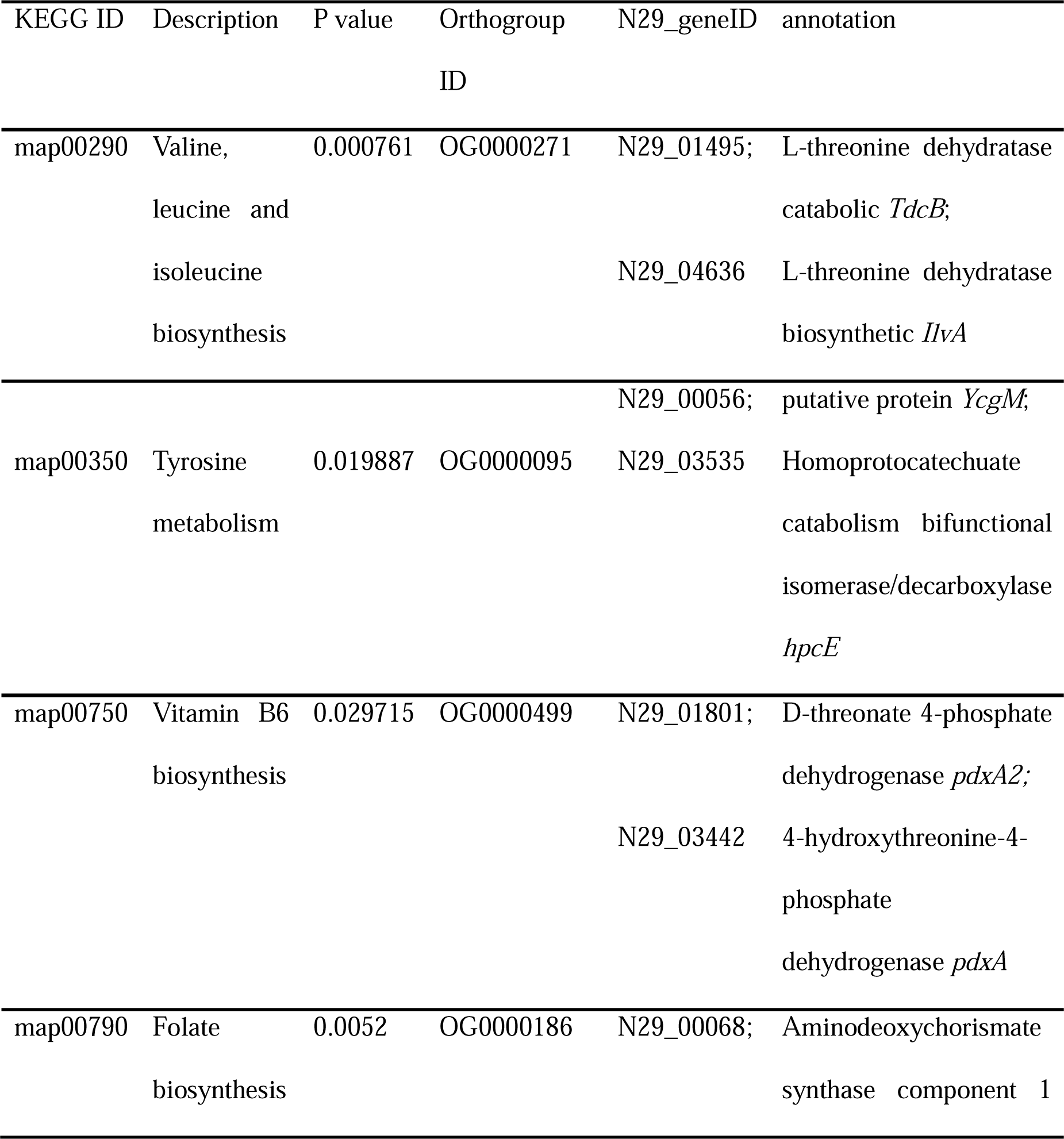
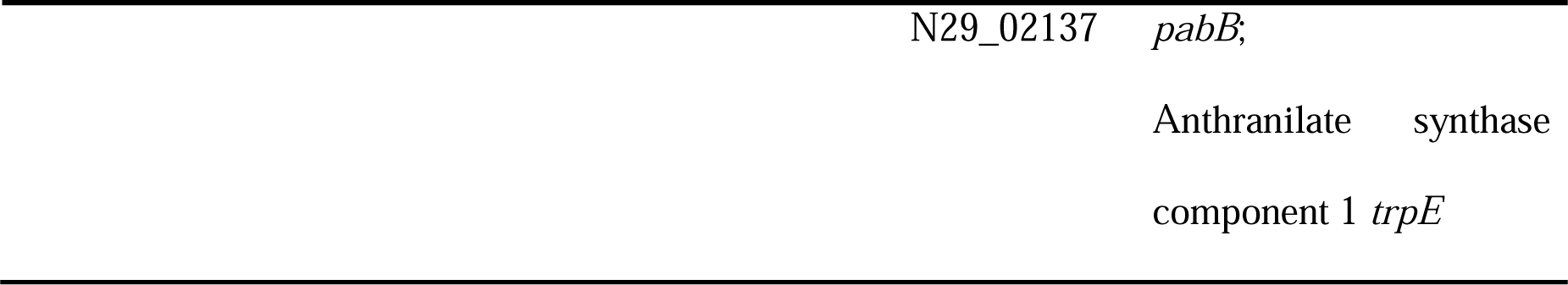
Candidate KEGG pathways and probiotic genes in N29 strain.

Further, we explored whether *E. cloacae* N29 could promote larval growth by producing a vitamin B6 (Figure 5B). Moreover, the growth of *E. cloacae* N29 and ΔpdxA mutant was similar in LB medium, indicating that the inability of the ΔpdxA mutant to promote larval size was not due to its own growth defect (Figure 5C). Then, we tried rescuing the phenotype of larval boby length by co-feeding *B. dorsalis* axenic larvae with *E. cloacae* N29 ΔpdxA mutant and vitamin B6. The results showed that the addition of vitamin B6 successfully restored *E. cloacae* N29 ΔpdxA mutant ability to rescue larval body length, indicating *E. cloacae* N29 promoted larval growth via enhancing vitamin B6 synthesis by *pdxA* (Figure 5D).

## Discussion

Here we found that the intestinal commensal bacteria promoted *B. dorsalis* larval growth under poor nutrition conditions, and this promotion effect was not limited to a single bacteria species. Our GWAS analysis revealed that microbiota genes positively associated with host larval growth were enriched in several physiological pathways. We further demonstrated that *E. cloacae* N29 promoted host larval growth by regulating *pdxA-mediated* vitamin B6 biosynthesis. In summary, our data reveal the important role of the gut microbiota in larval growth of *B. dorsalis*.

We found that *B. dorsalis* larvae harbored a large number of stage-specific low-abundance microbial communities. In contrast, high-abundance microbial communities were relatively stable in different larval stages. Furthermore, the richness and diversity of gut microbial communities were gradually decreased with larval development, which was in line with the previous report on *Bactrocera minax* [40]. Our data showed that Enterobacteriaceae and Leuconostocaceae were the most dominant species present in the *B. dorsalis* larval gut. Consistently, one previous study has also found that *Enterobacteriaceae* is the dominant bacterial species in the *B. dorsalis* larval gut in different geographic populations and under different rearing conditions [41]. In *B. minax*, the increase in the abundance of Leuconostocaceae during development has been reported to be associated with fructose and mannose metabolic activities in the intestinal tract [40]. In many insects, *Enterobacteriaceae* is a common bacterial species affecting host metabolism, immunity, and reproduction [42–47], based on which, we speculated that it might play an important role in *B. dorsalis* larval development.

We found that multiple microbial species rather than a single species were required for larval growth on a poor diet. Indeed, many studies have shown that microbiota could facilitate insect growth under nutrient-deficient conditions. Germ-free *Drosophila* larvae grow more slowly with smaller size on a poor diet. Mono-association with either one of two commensal bacteria, *Acetobacter pomorum* or *Lactobacillus plantarum* promotes larval growth through TOR-pathway and insulin pathway [32,33,48]. It has been widely reported that the microbiota produces numerous growth-promoting factors in *Drosophila* model [31,34,49,50]. Similar phenotypes were also observed in other insects such as *A. mellifera* [51] and *Ceratitis capitate* [52]. In the beetle *Holotrichia parallela*, microbiota participate in host metabolism and provide nutrients for host by degrading cellulose and hemicellulose [53–56]. Our previous study has reported that gut fungi also have a probiotic function in *B. dorsalis* [57]. However, the existing studies focused on only a few microbiota species and their functions in larval development. In this study, we found that at least over a dozen microbiota species could promote larval development of *B. dorsalis* Our GWAS results also suggested that multiple bacteria physiological pathways might be responsible for host development under nutrient deficiency conditions. For example, two biofilm formation pathways listed in the top five enriched pathways were positively associated with larval growth, suggesting microbiota biofilm formation could be the key factor regulating this host-microbe relationship in *B. dorsalis*. Actually, several studies have confirmed the necessity of biofilm formation in gut microbiota colonization [58–62]. Interestingly, a recent study has reported that outer membrane vesicle (OMV) biogenesis and biofilm-like aggregates enhance mosquito commensal bacterium colonization and the resistance of host to *Plasmodium* [63].

B vitamins are one of growth-promoting factors critical to the main metabolism of animals [49]. However, vitamin B cannot be synthesized by animals, and it is obtained from the environment or synthesized by the microbiota [64]. The microbiota in *Drosophila* could provide larvae with riboflavin, thiamine, and pantothenate to promote growth [34,50,65,66]. In addition, riboflavin biosynthesis is an important function for the gut microbiota in *A. aegypti* larvae [67]. Vitamin B provided by microbiota is involved in promoting *Glossina* development and reproduction [68]. Our study revealed that vitamin B6 biosynthesis by microbiota was essential for promoting larval growth of *B. dorsalis*. Gene *pdxA* is mainly responsible for biosynthesis of vitamin B6. The *pdxA*-deleted *E. cloacae* N29 mutant could not biosynthesize vitamin B6, and thus this mutant lost host larval growth promotion capacity. Vitamin B6 exists in six main forms, and pyridoxal-5’-phosphate, as one of main active forms of vitamin B6, is an important cofactor for more than 140 enzymes [68]. Similar to our observations, many bacteria such as *A. pomorum*, *L. plantarum*, *E. coli*, and *Rhodococcus rhodnii* have been predicted to have a complete vitamin B6 biosynthesis pathway in their genomes [68]. In tsetse fly, vitamin B6 produced by its symbiotic bacteria *Wigglesworthia glossinidia* is important for maintaining host proline homeostasis and fecundity [69]. In other insects such as aphids, bedbugs, kissing bugs, and ticks, diet supplementation of vitamin B6 partially rescued phenotype defect of larval development [68]. In this study, we found that of 15 strains promoting larval growth, *E. cloacae* N29 exhibited the most prominent promoting effect on larval growth of *B. dorsalis*. We speculated that growth-promoting effects of the 14 other bacteria species could also rely on vitamin B6 synthesis, suggesting a potential redundancy role of microbiota in supporting larval growth.

In summary, our data demonstrate that multiple microbiota species can promote larval development of the fruit fly *B. dorsalis,* and reveal the key role of vitamin B6 biosynthesis in promoting *B. dorsalis* larval growth. KEGG analysis indicates that multiple pathways, especially amino acid, lipopolysaccharide, fatty acid, and oxidative phosphorylation pathways, potentially promote larval growth of *B. dorsalis*. Our work paved the way for further functional studies focusing on insect-microbe relations under a more complex scenario, with multiple bacteria species-host interactions.

## Acknowledgments

This study was supported by the National Natural Science Foundation of China (No. U21A20222, No. 32220103009), National Key R&D Program of China (No. 2021YFC2600400), China Agriculture Research System of MOF and MARA (CARS-26) and Hubei Hongshan Laboratory.

**Availability of supporting data**

**Sequence files and metadata for all samples used in this study and all data supporting the main conclusions have been deposited in Figshare (DOI: 10.6084/m9.figshare.25039781.**

**Private link:**https://figshare.com/s/6c0e87678053eb95f6f9

## Materials and methods

### Insect rearing

The experimental insects were collected from Guangdong Province, China, using protein bait and maintained in the Institute of Urban and Horticultural Insects, Huazhong Agricultural University, Wuhan, Hubei, China. The photoperiod of the insect-rearing room was 12 h light:12 h dark. The room’s relative humidity was 70– 80%, and the temperature was maintained at 28 ± 1[. Larvae were fed with larval food (wheat bran, 80 g; corn flour, 40 g; sucrose, 40 g; yeast powder, 15 g; water, 200 mL). After eclosion, adult flies were moved to 30 cm × 30 cm × 30 cm cages. Adult flies were raised on a sucrose and yeast mix at a ratio of 3:1.

### Preparation of larval gut samples

The larvae were rinsed in 75% alcohol for 3 min, followed by three rinses in sterile water. An appropriate amount of sterile PBS buffer was added to a sterile Petri dish. The larvae were dissected with sterile forceps under a stereopicroscope to obtain gut samples. The resultant gut samples were used for total DNA extraction and the detection and isolation of culturable bacteria. There were 5 biological replicates for each instar larval gut sample, with 30 larval gut tissues per biological replicate.

### Bacterial DNA extraction and 16S rDNA amplicon sequencing

Bacterial DNA was extracted from each instar larval gut sample using E.Z.N.A.® Soil DNA Kit (Omega, Norcross, GA, USA), according to the manufacturer’s instructions, with five biological replicates. The variable region V3+V4 of the 16S rRNA gene was amplified using a broad-range primer pair (338F: 5’-ACTCCTACGGGAGGCAGCA-3′, 806R:5′-GGACTACHVGGGTWTCTAAT-3′) using the Phusionâ High-Fidelity PCR Master Mix (New England Biolabs, Beverley, MA). The PCR amplification program was as follows: preincubation at 95□ 5 min; followed by 35 cycles of at 56□ 45 s, at 72□ 1 min, and at 94□ 45 s, ending up with a final extension at 72□ 10 min. The PCR products were subjected to PE300 double-end sequencing on the Illumina MiSeq sequencing platform.

### Analysis of 16S rRNA gene amplicon data

In order to integrate the raw double-end sequencing data, FASTQ double-end sequences were filtered using the sliding window method, and parameters were set as follows: The window size was 10 bp; the step size was 1 bp; the average quality score of the sequence in the window was ≥ Q20; and the average sequencing accuracy of the base was ≥ 99%. The sequence whose average quality score was lower than Q20 was truncated from the first window, and the truncated sequence length was ≥ 150 bp. The ambiguous base N was not allowed. After preliminary quality control, the paired-end sequences were pairwise connected based on overlapping bases using FLASH software following the requirements that the overlapping base length between read 1 and read 2 was ≥ 10 bp, and that base mismatches were not allowed. Finally, valid sequences for each sample were obtained. Chimeric sequences were removed using USEARCH. The resultant qualified sequences were merged and divided into different (operational taxonomic units) OTUs based on 97% sequence similarity using QIIME software. The sequence with the highest abundance in each OTU was selected as the representative sequence of this OTU. Subsequently, based on the number of sequences contained in each OTU in each sample, a matrix file of the OTU abundance in each sample was constructed. By aligning the OTU representative sequence with the sequence in the Greengenes database, the taxonomic information of the corresponding OTU was obtained. Diversity indexes such as Simpson index, Chao1 index, ACE index, and Shannon index were calculated for each sample using QIIME software. The relative abundance matrix at the genus level was submitted to the Galaxy online analysis platform for LEfSe analysis.

### Isolation and identification of culturable bacteria from larval gut

Each instar larvae gut tissue samples were put into a 1.5ml centrifuge tube and added with 1 ml sterile PBS and an appropriate amount of sterile glass sand. The 10 larval gut tissue samples of each instar was homogenized, and then 100 μl of the homogenate was gradient diluted (by10×, 10^-4^, 10^-5^, and 10^-6^). The 100 μl of homogenate was coated onto nutrient agar (NA) medium and violet red bile glucose agar (VRBG) medium and cultured at 30□ for 24 −48 h under aerobic conditions, or incubated on DeMan-Rogosa-Sharpe (MRS) medium at 30□ for 24-48 h in an MGC-7L-sealed anaerobic incubator. Next, 100 single colonies were randomly selected from each of the three medium plates of gut tissue homogenate culture at 3 different instars (1st-3rd) and placed in the corresponding liquid media (NA, VRBG, and MRS). The colonies were incubated in NA and VRBG liquid media at 200 r/min for 24 h at 30□, whereas the colonies were incubated in MRS liquid media were incubated anaerobically for 24 h at 30 ° C. After the incubation, the resultant bacterial solution was added with an equal volume of 40% glycerol and stored in the refrigerator at −80□. Bacterial genomic DNA was extracted from the bacterial solution using HiPure Bacterial DNA kit (Magen, Guangzhou, China).

### Mono-association experiment between bacteria and larvae

Referring to the axenic drosophila construction method, bacteria and larvae were cultured. The specific procedures were as follows:

Preparation of related strains. Single colonies of isolated and identified strains were put into corresponding liquid media (NA, VRBG, and MRS) and cultured under appropriate conditions.

Quantification of bacteria in bacterial solution. After the bacterial solution was diluted with the liquid medium at the ratio of 1:1, 1:2, and 1:4, OD600value was measured. These diluted bacterial solution was coated onto the solid medium and cultured for 24 hour, and colony forming units (CFU) were counted. Sterile water volume for preparing bacterial resuspension solution was calculated according to the following formula: E = ((O-B) x V x D)/C. Where E is the volume of sterile water required for preparing bacterial resuspension solution; O is the OD600 value of cultured bacterial solution; B is the OD600 value of blank culture solution; D is the dilution folds; V is the volume of bacteria suspension collected by centrifugation, and the above volume unit was expressed as microliters (μl).

Incubation of bacteria and larvae. The 100μl bacterial solution was added to the sterile tube containing irradiation-sterilized larval feed and sterilized eggs and incubated in a constant temperature incubator at 27□ and a humidity of 70% for 6 days. (4) Determination of larval growth and development. Specifically, the feeding tube was opened, and the feed and larvae were poured onto the petri dish. The larvae were collected into the EP tube containing 50% glycerol with forceps. Subsequently, the larvae were placed on ice for 30 min. The photos of larvae were taken with the Olympus stereoscope. The larval body length was measured.

### Next-generation library construction and sequencing

Quality control of bacterial genomic DNA was performed as follows. Agarose gel electrophoresis was used to analyze the purity and integrity of DNA, and NanoDrop 2000 Spectrophotometer (Thermo Fisher Scientific Inc.) was used to detect the purity of DNA (OD 260/280 ratio). Qubit 2.0 was used to determine the DNA concentration. After quality control, the bacterial genomic DNA samples were randomly interrupted by the Covaris ultrasonic breaker. Afterwards, a 350 bp library was constructed by the following steps: end repair, addition of A-tail, addition of sequencing adapter, purification, and PCR amplification. Subsequently, library concentration was measured using Qubit 2.0 and diluted, and the fragments inserted into library were detected using Agilent 2100. After the size of the insert fragment met the requirement, the effective concentration of the library was quantified by the q-PCR method to ensure the quality of the library. After the library quality was qualified, library was subjected to PE150 sequencing on the Illumina Novaseq 6000 high-throughput sequencing platform.

### De novo genome assembly and annotation

The main steps for de novo genome assembly and annotation were as follows: Illumina paired-end data were corrected and assembled using SPAdes software. The numbers of contigs and dead ends were required to be minimized after assembly. The contigs were further constructed into a genome scaffold diagram. According to the depth of reads and the connectivity between the assembly diagrams, single-copy contigs were distinguished from multi-copy contigs using the Unicycler software by the greedy algorithm. The final genome assembly results for each strain were evaluated using QUAST software, in terms of the number and length of contigs, the maximum contig number, and the N50 value in the assembled genome. The coding sequences, ribosomal RNA genes, transfer RNA genes, signal peptide genes, and non-coding RNA genes in the genome were annotated using Prokka software.

### Whole-genome phylogenetic analysis

The fasta files of 27 strain genomes were converted into Anvi’o files, which were further converted into a single Anvi’o contig database. Subsequently, a single-copy gene HMM file was constructed, and the single-copy gene amino acid sequences of the 27 genomes were extracted based on the HMM file. These sequences were concatenated and aligned to generate a fasta file. The aligned single-copy gene amino acid sequences were used to construct a phylogenetic tree.

### Whole-genome association analysis of bacteria

The genomes of larval intestinal strains were subjected to second-generation high-throughput sequencing, assembly, and annotation, to obtain orthologous genes of 27 strain genomes. The statistical analyses of orthologous genes present in or absent from different strains and the larval body length data were performed using MAGNAMWAR in R package. The orthologous genes significantly associated with the phenotype were screened by calculating the association between the phenotype and the orthologous genes, with p < 0.05 as screening threshold, to obtain an orthologous gene set. The orthologous gene sequences in all larval intestinal strain genomes were functionally annotated using eggNOG-mapper software based on EggNOG 5.0 Database to obtain the GO and KEGG annotation information of orthologous genes. GO and KEGG enrichment analyses of orthologous gene sets that were significantly related to larval body length were performed using clusterProfiler in R package, with p < 0.05 as enrichment analysis screening threshold.

### Construction of bacterial mutant strains

We used the λ-Red homologous recombination system to knock out candidate bacterial genes. The specific procedures were as follows:

Preparation of N29 electroporation competent cells. The N29 strain was cultivated at 37°C at 200 rpm/min until OD600 reached 0.4 - 0.6. The bacterial solution was centrifuged at low temperature for 15 min. The strains were collected, resuspended in 30 mL of pre-cooled 10% glycerol, and stored in a −80°C refrigerator. The 50 ng of pKD46 plasmid was added to 100 μL of N29 electroporaion competent cells, and mixed gently, and the mixture was transferred to a pre-cooled electroshock cup. The electroshock cup was placed in the bio-rad electroporator for electroporation with voltage set as 1.8 kV. Then 1 ml of LB was added, followed by shaking culture and recovery. The bacterial solution was coated onto LB solid medium containing ampicillin and cultured at 30°C overnight. The single colonies were collected from solid LB medium containing ampicillin and cultured in LB liquid medium with shaking at 30°C overnight. Lambda red induction reagent was added at a ratio of 1:200. Bacterial solution was cultured with shaking until OD600 reached proximately 0.4 - 0.6. Then bacterial solution was placed on ice for 30 min and centrifuged at low temperature for 15 min. The strains were collected, resuspended in 30 mL of pre-cooled 10% glycerol, and stored in −80□ refrigerator. A 70 bp homologous recombination primer pair was designed. Of 70bp, 50 bp of the upstream and downstream primer pair was homologous sequence complimentary to the gene to be knocked out, and the remaining 20 bp was complementary to the sequence of pKD4. Then 500 ng of PCR product was added to 100 μl of electroporation competent cells and mixed gently, and the mixture was transferred to a pre-cooled electroshock cup. The electroshock cup was put into the bio-rad electroporator for electroporation, with voltage set as 1.8 kV. The 1 ml of LB was added for shaking culture and recovery, and then bacterial solution was coated onto LB solid medium containing kanamycin, and cultured at 37°C overnight. The genome of the mutant strain was extracted and PCR amplified using 20 bp primers (Table S2).

### Vitamin B6 content determination

The 1ml bacterial culture medium was added into in 1.5ml EP tube and centrifuged at 13000 g for 10min. The 40μl supernatant was collected for vitamin B6 content determination using vitamin B6 content kit (Boxbio, Beijing, China) according to the manufacturer’s instructions.

### Fed Vitamin B6 assay

The 200ul 5mg/ml vitamin B6 solution (Sangon Biotech, Shanghai, China) and pdxA-deleted N29 mutant strain were co-fed to the larvae.

### Data statistics and analysis

GraphPad Prism 9.0 software and R (3.6.3) were used for statistical analysis and visualization of the data. The differences between two independent samples were analyzed using parametric Student’s T test. Multiple comparisons of multiple samples were performed by Tukey’s multiple comparison method in one-way ANOVA, and P <0.05 was considered as statistically significant.

---

**Figure S1.**
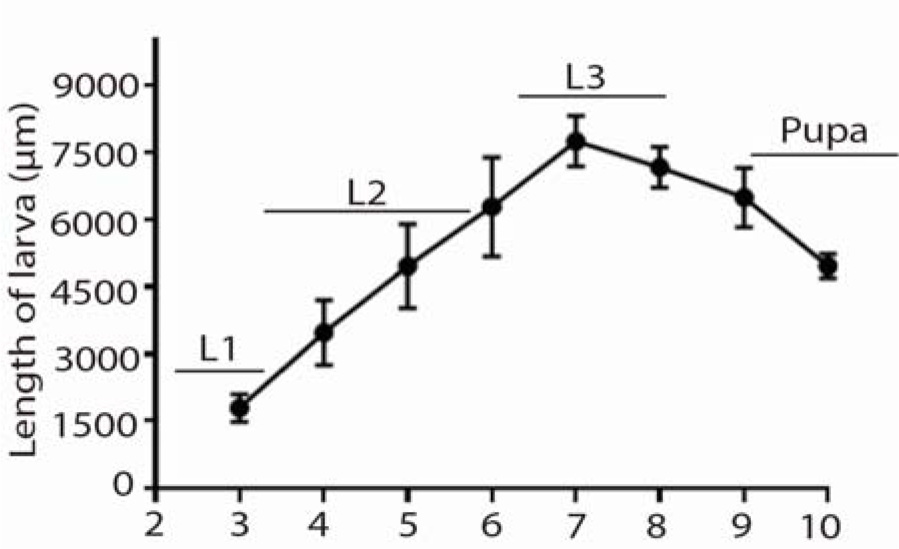
The growth of larvae fed on bananas diet. The body length of the larva from day 3 to day 10 after laying eggs. The data point represent the length of at least 20 larvae (mean + SE). Length unit: μm. L1: First instar larva; L2: Second instar larva; L3: Third instar larva.

**Table S1.**
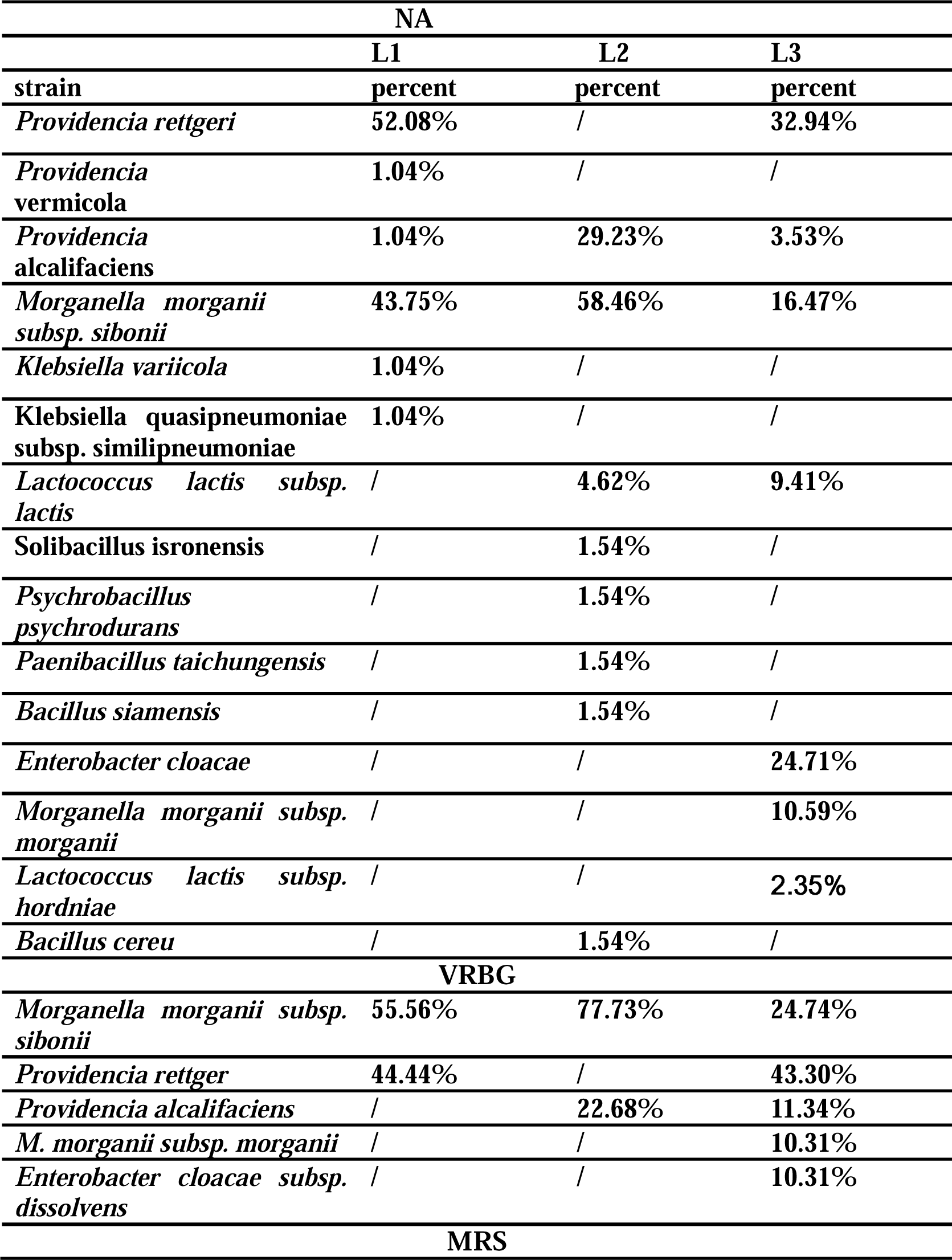
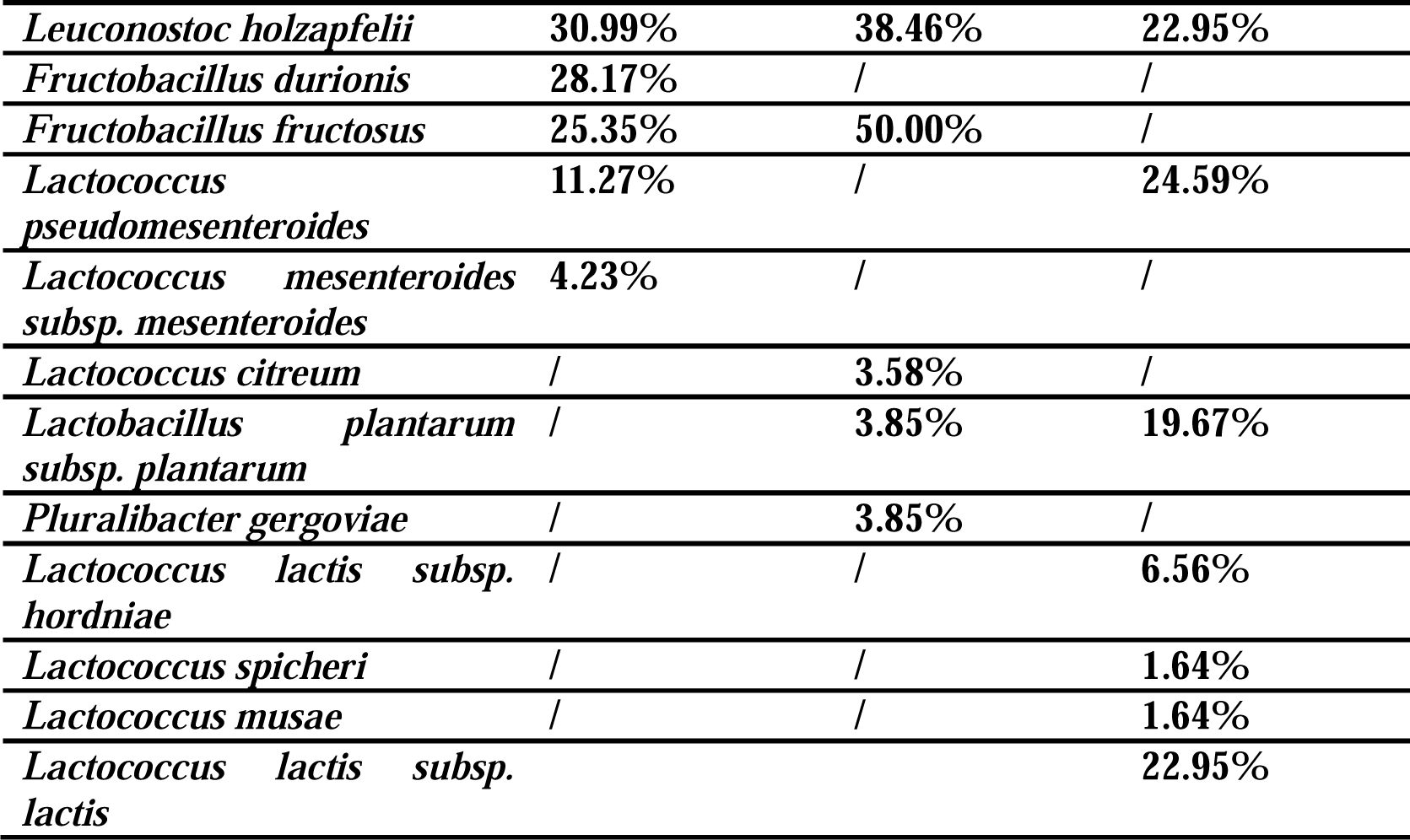
Proportion of intestinal microbiota of different instars in the three media.

**Table S2.**
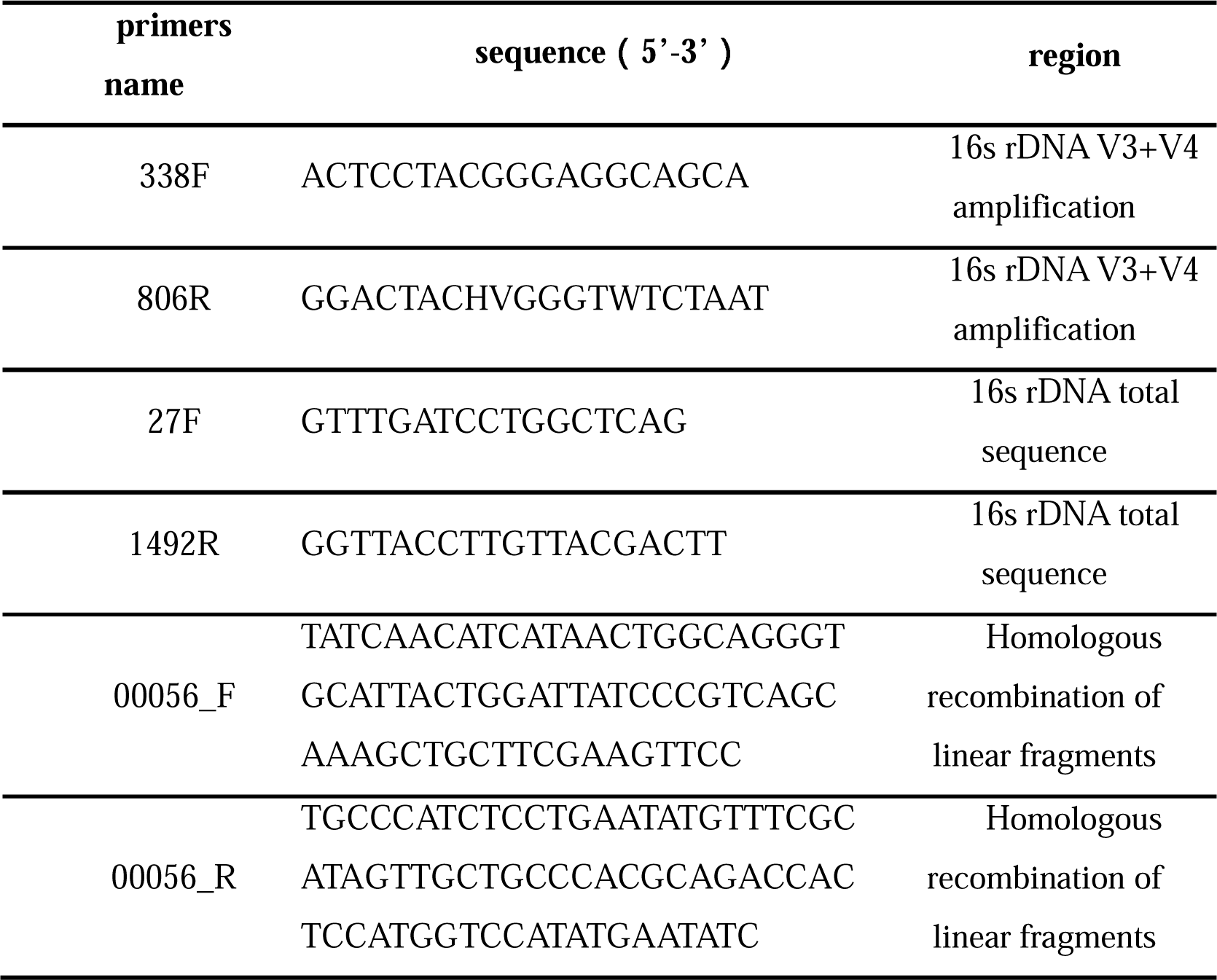
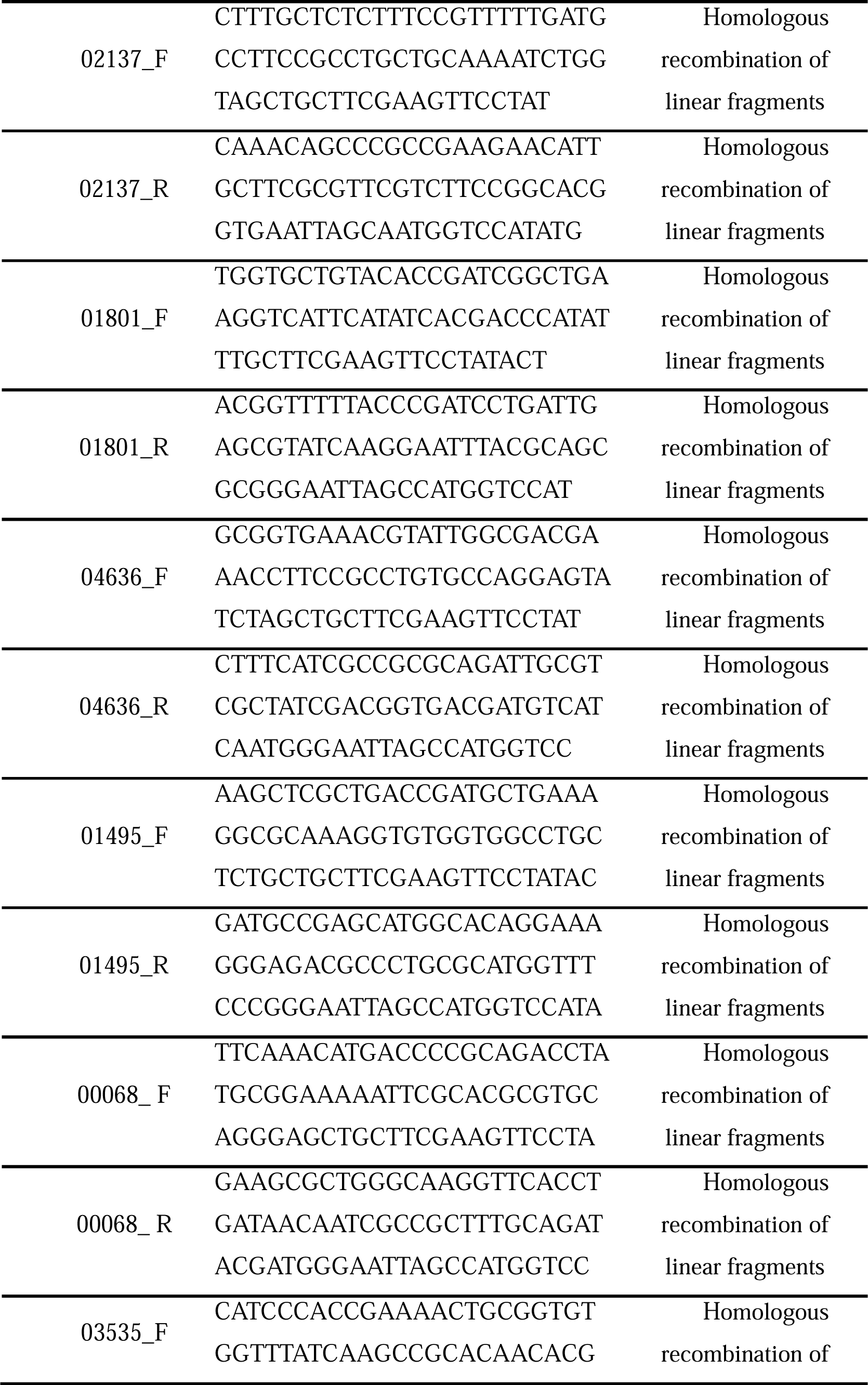
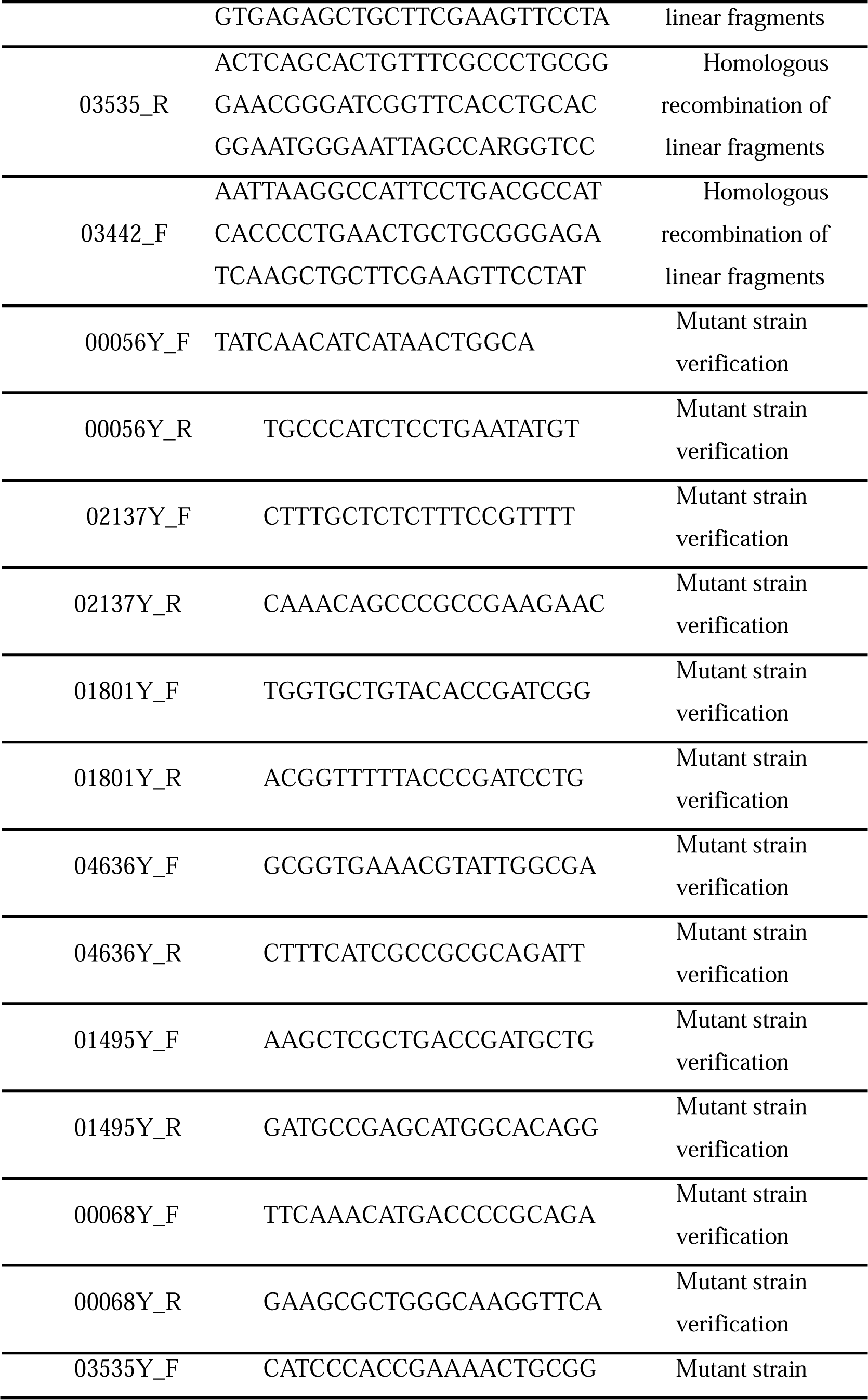
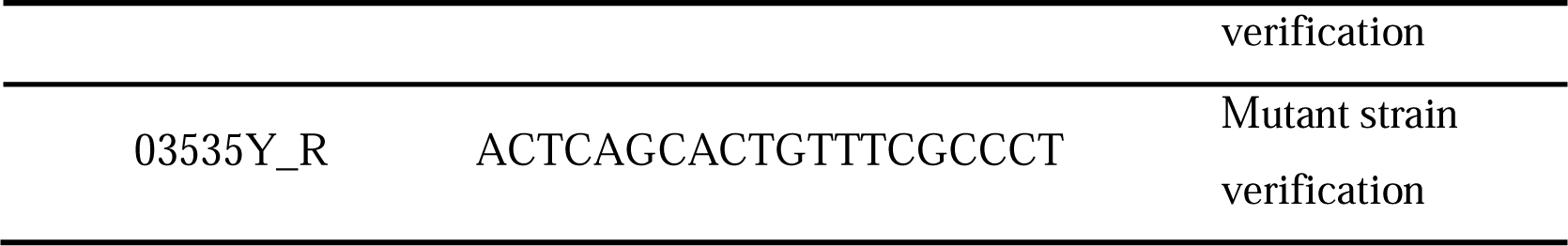
The primers used in the λ-red homologous recombination primers.

